# Coding nucleic acid sequences with graph convolutional network

**DOI:** 10.1101/2022.08.22.504727

**Authors:** Ruo Han Wang, Yen Kaow Ng, Xianglilan Zhang, Jianping Wang, Shuai Cheng Li

## Abstract

Genome sequencing technologies reveal a huge amount of genomic sequences. Neural network-based methods can be prime candidates for retrieving insights from these sequences because of their applicability to large and diverse datasets.However, the highly variable lengths of nucleic acid sequences severely impair the presentation of sequences as input to the neural network. Genetic variations further complicate tasks that involve sequence comparison or alignment. Here, we propose a graph representation of nucleic acid sequences called *gapped pattern graphs*. These graphs can be transformed through a Graph Convolutional Network to form lower-dimensional embeddings for downstream tasks. On the basis of the gapped pattern graphs, we implemented a neural network model and demonstrated its performance in studying phage sequences. We compared our model with equivalent models based on other forms of input in performing four tasks related to nucleic acid sequences—phage and ICE discrimination, phage integration site prediction, lifestyle prediction, and host prediction. Other state-of-the-art tools were also compared, where available. Our method consistently outperformed all the other methods in various metrics on all four tasks. In addition, our model was able to identify distinct gapped pattern signatures from the sequences.

## Introduction

The advances in next-generation sequences technology lead to explosive accumulation of nucleic acid sequences. Accurate characterization of those newly discovered sequences, which is inherently a problem of labeling new sequences with the knowledge from known sequences, improves our understanding of biological systems. Thus, there is a critical need for computational tools to analyze nucleic acid sequences.

The homology search serves as a primary approach to studying sequences. Recently, homology has been computed with alignment-free (AF) methods to avoid the pitfalls of sequence alignment^1^. These AF methods calculate the frequency or uniqueness of different *k*-mers to characterize genomes, and use various statistical measures to determine the similarity between different genomes. However, several drawbacks limit the robustness of these AF approaches. First, approaches based solely on *k*-mer distributions forfeit all possibilities of capturing crucial information on the interaction between *k*-mers or the order of their occurrences. Second, most AF approaches require exact matches of *k*-mers, which overlook the frequent genetic variations in biological sequences, including single nucleotide polymorphisms (SNPs) and insertions or deletions (InDel).

An alternative to AF-based inputs is one-hot encoding of sequences, which are commonly used in neural network models for bio-sequence analyses. However, such encodings adapt poorly to sequences of variable length. They also have no capability to associate neighboring nucleotides into frequently occurring components^2^.

An additional drawback restricts all such vector-based (or matrix-based) inputs, whether one-hot or AF, since they inevitably introduce an ordering over the encoded features. For instance, a vector of *k*-mers would implicitly identify each *k*-mer by a chosen position within the vector. An arbitrarily chosen ordering would result in biases in the trained neural network models. On the other hand, nonarbitrary ordering is hard to realize. For instance, one can require that neighboring elements of the vector represent *k*-mers that differ only by a single nucleotide. However, since each *k*-mer has 3*k* neighbors that differ by a single letter, this ordering cannot be performed with vectors (or matrices).

A simpler and more natural encoding is to represent each *k*-mer as a vertex in a graph and model the relationships (or interactions) between *k*-mers as graph edges. This way, there would be no implicit ordering on the *k*-mers. We call such a graph a *gapped pattern graph* (GPG), and construct it as follows. Given a sequence, we create a vertex to represent each *k*-mer in the sequence; the vertex is assigned a feature that encodes the frequencies of the *k*-mers within the sequence. An edge connects two vertices if their corresponding *k*-mers appear within close proximity at least once in the sequence. The construction hence (1) encodes the *k*-mer distribution; and (2) enables a walk of the graph to represent a potential pattern (in the form of a few *k*-mers with gaps in-between) within the sequence. These *k*-mers with gaps attempt to resemble the patterns that contain *spaced seeds*^3^, which are known to tolerate minor substitutions, InDels, sequence errors, *etc.*.

The comparison of two GPGs can give us the similarity between the sequences they represent. Such comparisons can be made naturally through so-called graph neural networks^4^, which have been studied very intensely in recent years. We train a Graph Convolutional Network (GCN)^5^ that transforms each GPG into a vector in a low-dimensional latent space; the vectors are then used in downstream analysis tasks. We call our resultant framework *gapped pattern-GCN* (GP-GCN). To our knowledge, the framework is the first method that utilizes GCN to explore pattern relationships in biological sequences.

In this study, we demonstrated the superiority of the GP-GCN framework by applying it to phage sequences (i.e., virus-infecting bacteria or archaea). The length and genomic variations of the phage sequences make their analysis a challenging task. Under the aegis of the GP-GCN framework, we developed Graphage, a tool that performs four phage-related tasks: phage and ICE discrimination, phage integration site prediction, phage lifestyle prediction, and phage host prediction. Graphage achieves state-of-the-art performance in tasks and mines distinct gapped pattern signatures for phage phenotypes.

## Results

### Overview of GP-GCN framework and Graphage

In this study we present the GP-GCN framework to encode nucleic acid sequences. In the input preparation stage, each phage sequence *S* is converted into a GPG (see *Gapped Pattern Graph Construction* subsection). Each of the sequence’s constituent *k*-mers becomes a vertex, and is assigned the *k*-mer’s frequency as its feature. An edge points from a vertex of the *k*-mer s to a vertex of the *k*-mer *t* if and only if *t* occurs after s at least once in *S* within a given threshold distance. Therefore, such an edge represents a pattern of the form *sxt*, where *s* and *t* are *k*-mers and *x* is an arbitrary string of length up to the given threshold. The edge is assigned the frequency of the pattern *sxt* in *S* as its feature.

The first part of GP-GCN transforms each GPG into a low-dimensional embedding (latent space vectors) using a multilayer GCN (see *Neural Network Architecture* subsection) (Figure 1A). The second part implements fully-connected neural network models for prediction tasks (Figure 1B). These models accept embeddings as input and give predictions as output for a specific task. The GP-GCN framework features a modular design that can be flexibly applied to genomic study. The GP-GCN framework is available at https://github.com/deepomicslab/GCNFrame.

**Figure 1.**
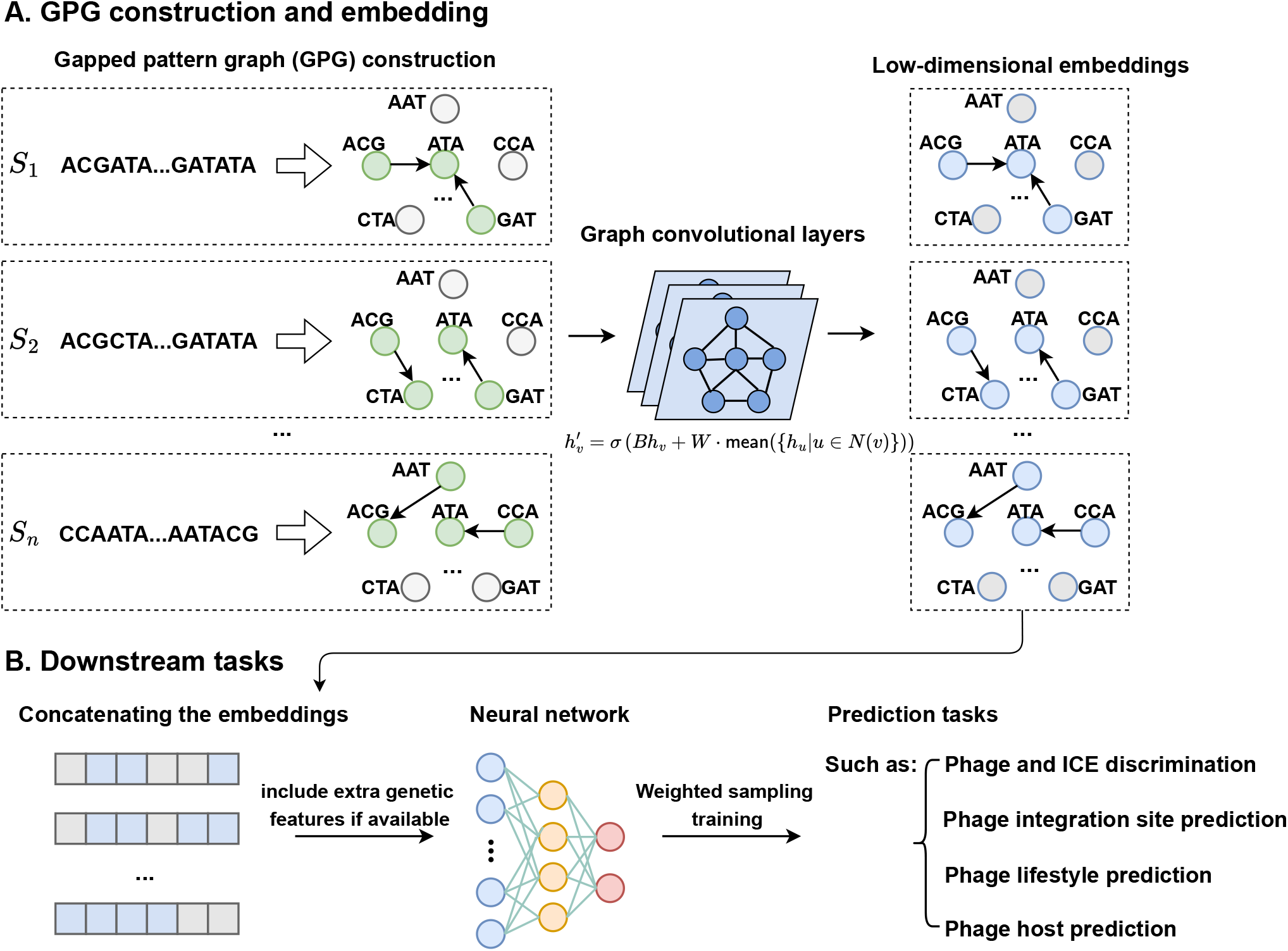
Overview of GP-GCN framework. A. A gapped pattern graph is constructed for each sequence and is embedded into latent spaces with a GCN. B. The embeddings are given as input to neural networks for downstream tasks.

Applying the GP-GCN framework to phage sequences, we designed Graphage for four phage-related tasks. Four neural network models are used, each corresponding to a specific phage analysis task. The GCN and the downstream model are trained simultaneously. Graphage is available at https://github.com/deepomicslab/Graphage.

### Phage and ICE discrimination

We use the sequences from NCBI^6^ and ICE^7^ as the training set for phage and ICE discrimination. A classifier is used to distinguish between the phage and ICE sequences given their respective GP-GCN embeddings. Following the practice introduced in GPD^8^, we include information on gene density and hypothetical protein fraction as input to the classifier in addition to embeddings. We optimize two hyperparameters by grid search: (1) the maximum allowed length *d* for the gap between *k*-mers, and (2) the number of layers *l* in the GCN. The results showed that *d* = 1 and *l* = 3 achieved the best performance (Supplementary Figure S1). More hyperparameter settings can be found in Supplementary Table S1.

We first compared the effects of using the GP-GCN embeddings with those of using other input types. When tested in the ImmeDB database^9^, Graphage outperformed AF-based, word2vec-based, and sequence-based models, with a 7.26% increase in average accuracy, 0.1117 increase in F1-score, and 0.0181 increase in average area under ROC (AUC), compared to the respective second-highest scores (Table 1). The sequence-based model gave the worst performance, consistent with our argument that the method is unsuitable for sequences of variable lengths.

**Table 1.**
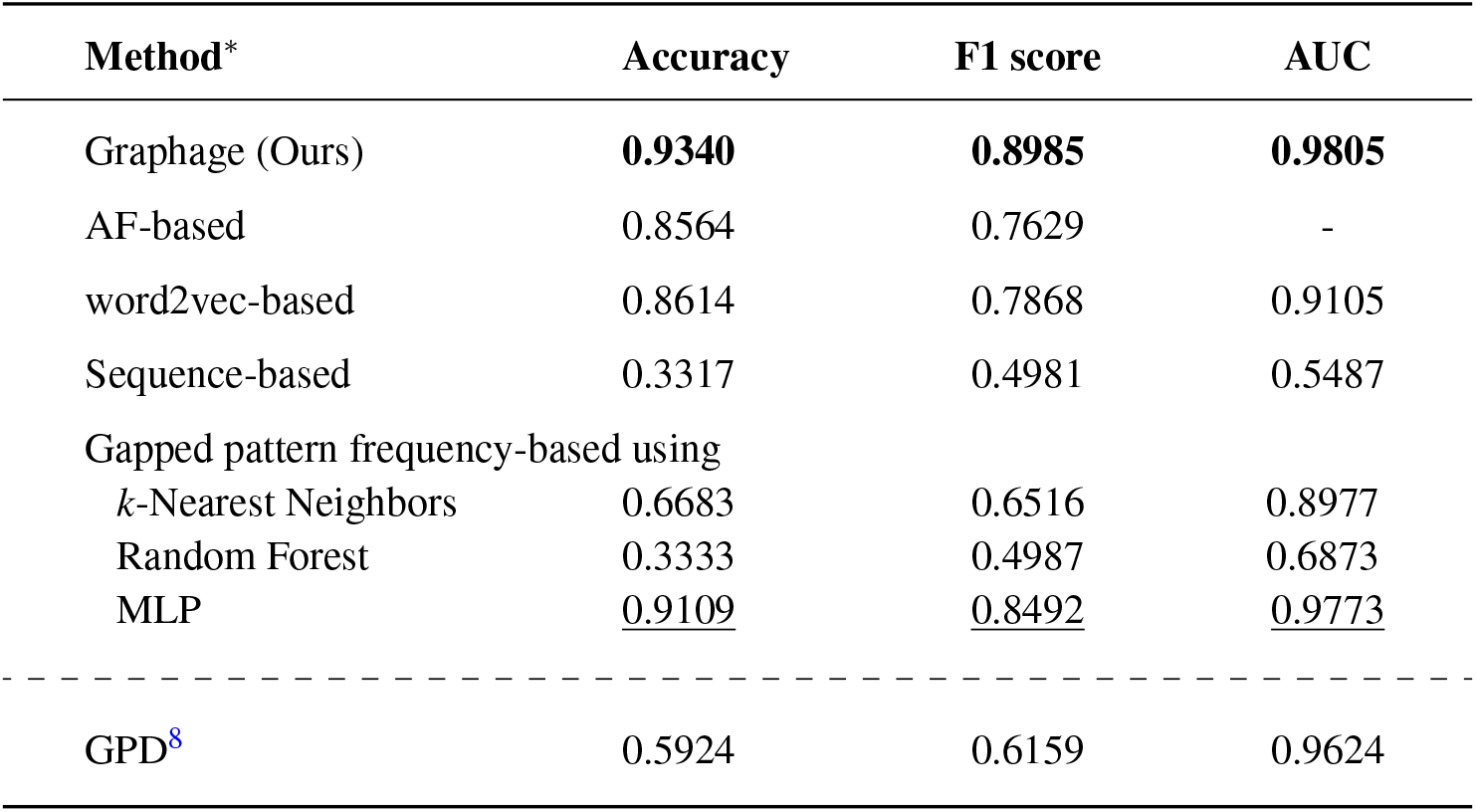
Comparing Graphage with other models/tools on phage and ICE discrimination tasks. The best-performing model is in bold font, while the runner-up is underlined. AUC is inapplicable for the AF-based model since the model does not give probabilities.

| Method* | Accuracy | F1 score | AUC |
| --- | --- | --- | --- |
| Graphage (Ours) | **0.9340** | **0.8985** | **0.9805** |
| AF-based | 0.8564 | 0.7629 | - |
| word2vec-based | 0.8614 | 0.7868 | 0.9105 |
| Sequence-based | 0.3317 | 0.4981 | 0.5487 |
| Gapped pattern frequency-based using |  |  |  |
| *k*-Nearest Neighbors | 0.6683 | 0.6516 | 0.8977 |
| Random Forest | 0.3333 | 0.4987 | 0.6873 |
| MLP | <u>0.9109</u> | <u>0.8492</u> | <u>0.9773</u> |
| ----- |  |  |  |
| GPD<sup>8</sup> | 0.5924 | 0.6159 | 0.9624 |

Since a GPG contains information on frequencies of both the *k*-mers (vertex feature) and the gapped patterns of two *k*-mers (edge feature), it is worth questioning if the latter could be solely responsible for the performance of Graphage. To answer this question, we performed tests with the frequencies of the gapped patterns as input to several conventional machine learning models (see Competing Methods subsection). The results show that this is not the case (Table 1).

We then compared Graphage to the model introduced in GPD^8^ for phage and ICE discrimination. We found that the GPD model is prone to misclassify ICEs as phages, leading to lower accuracy and F1 scores.

Finally, there is the concern of whether Graphage’s classifier depended more on the embeddings or the genetic features (i.e., gene density and hypothetical protein fraction) for its output. To evaluate this possibility, we re-evaluated Graphage with only the embeddings or only the genetic features as input. We found that the best performance is achieved with both inputs, showing that they jointly contributed to the performance (Supplementary Figure S2).

### Phage integration site prediction

Phage integration site inference is complicated by the fact that circular phage genomes can integrate into the bacterial genome as prophages. To distinguish them, we trained two models that predict phage integration sites on phage genomes and bacterial genomes. A dataset is prepared for each model. The data set for phage integration site consists of 21,117 positive and 21,117 negative samples; that for bacterial integration site consists of 17,550 positive and 17,550 negative samples. We used 90% of the data as training and validation sets. In addition to the allowed gap length *d* and the number of graph convolutional layers *l*, we also optimized the window length *w* (i.e., the input sequence length). The results show that *d* = 2, *l* = 3, and *w* = 600 gave the best performance for integration site prediction for both models (Supplementary Figure S3 and Figure S4). More hyperparameter settings are shown in Supplementary Table S2.

When evaluated with the respective testing set, both models obtained prediction precisions greater than 0.8, significantly outperforming the models based on AF, word2vec, sequence, or gapped pattern frequencies under various metrics (Tables 2 and 3). For integration site prediction in phage sequence, Graphage has a 3.90% lead on average accuracy over the runner-up’s score; this lead is 2.80% with bacteria genomes. We observe that the sequence-based model performed worse when inputs are of variable length than when they are of the same length, demonstrating the disability of the representation in handling length variations.

**Table 2.** Comparing our graph model with other models/tools on phage integration site prediction tasks. The best performing model is in bold font, while the runner-up is underlined. Results are averaged over 10 random train-test splits.

| Method | Accuracy | F1 score | AUC |
| --- | --- | --- | --- |
| Graphage (Ours) | **0.849±0.005** | **0.839±0.006** | **0.882±0.004** |
| AF-based | <u>0.810±0.004</u> | <u>0.812±0.004</u> | - |
| word2vec | 0.791±0.007 | 0.779±0.010 | 0.844±0.008 |
| Sequence-based | 0.736±0.013 | 0.734±0.007 | 0.809±0.007 |
| Gapped pattern frequency-based using |  |  |  |
| *k*-Nearest Neighbors | 0.690±0.006 | 0.741±0.006 | 0.801±0.006 |
| Random Forest | 0.802±0.008 | 0.786±0.006 | <u>0.874±0.002</u> |
| MLP | 0.805±0.006 | 0.797±0.006 | 0.854±0.004 |

**Table 3.** Comparing our graph model with other models and tools on bacterial integration site prediction tasks. The best performing model is in bold font, while the runner-up is underlined. Results are averaged over 10 random train-test splits.

| Method | Accuracy | F1 score | AUC |
| --- | --- | --- | --- |
| Graphage (Ours) | **0.813±0.005** | **0.801±0.007** | **0.860±0.006** |
| AF-based | 0.772±0.007 | <u>0.782±0.008</u> | - |
| word2vec | 0.769±0.006 | 0.763±0.006 | 0.836±0.006 |
| Sequence-based | 0.735±0.010 | 0.732±0.012 | 0.820±0.012 |
| Gapped pattern frequency-based using |  |  |  |
| *k*-Nearest Neighbors | 0.588±0.031 | 0.690±0.006 | 0.681±0.008 |
| Random Forest | <u>0.785±0.021</u> | 0.767±0.031 | <u>0.855±0.007</u> |
| MLP | 0.774±0.006 | 0.769±0.007 | 0.833±0.007 |

### Phage lifestyle prediction

In lifestyle prediction, we want to classify phages by whether they are virulent or temperate. Since sequences alone are likely insufficient for this classification, we supplemented the classifier with additional information by using a dataset that contains 206 lysogeny-associated proteins as established by BACPHLIP^10^. We used HMMER^11^ to check the presence of the 206 proteins in each sequence and encoded the information into a 206-dimensional vector, with 0 for absence and 1 for presence. The maximum allowed gap length (*d*) and the number of graph convolutional layers (*l*) were optimized through grid search. The model with *d* = 2 and *l* = 4 achieved the best classification performance (Supplementary Figure S5). More hyperparameters can be found in Supplementary Table S3.

Graphage gave the highest average performance among all competing methods, including AF-based, word2vec-based, sequence-based, and gapped pattern frequencies-based models, as well as two phage lifestyle prediction tools, BACPHLI^P10^ and DeePhage^12^ (Table 4). Compared to the other methods, Graphage demonstrated large improvements in accuracy and F1-score metrics.

**Table 4.** Comparing our graph model with other models and tools on phage lifestyle prediction tasks. The best performing model is in bold font, while the runner-up is underlined. Results are averaged over 10 random train-test splits.

| Method | Accuracy | F1 score | AUC |
| --- | --- | --- | --- |
| Graphage (Ours) | **0.957±0.011** | **0.928±0.022** | **0.975±0.010** |
| AF-based | 0.894±0.014 | 0.836±0.310 | - |
| word2vec | <u>0.939±0.011</u> | <u>0.898±0.021</u> | 0.959±0.009 |
| Sequence-based | 0.810±0.047 | 0.657±0.064 | 0.858±0.023 |
| Gapped pattern frequency-based using |  |  |  |
| *k*-Nearest Neighbors | 0.875±0.023 | 0.803±0.044 | 0.926±0.016 |
| Random Forest | 0.890±0.018 | 0.805±0.040 | 0.948±0.019 |
| MLP | 0.938±0.017 | 0.895±0.035 | 0.967±0.011 |
| ----- |  |  |  |
| BACPHLIP<sup>10</sup> | 0.932±0.007 | 0.888±0.016 | 0.956±0.009 |
| DeePhage<sup>12</sup> | 0.845±0.030 | 0.875±0.024 | <u>0.969±0.010</u> |

To determine the contribution of the additional information, we trained two Graphage models, one with only embeddings and one with only information on lysogeny-associated proteins. Although these models performed worse than when both information were available (Supplementary Figure S6), the model with only protein information had an edge over that with only embeddings.

### Phage host prediction

Phage host prediction is a multiclass prediction problem. Our constructed dataset consists of 107 classes, each for a host species. We use 90% of the data for training and validation. Accuracy, weighted F1-score, and macro F1-score were used as evaluation metrics (see Methods). Under these metrics, the model with *d* = 2 and *l* = 1 achieved the best performance (Supplementary Figure S7). More hyperparameter settings are shown in Supplementary Table S4.

In addition to AF-based, word2vec-based, sequence-based, and gapped pattern frequencies-based models, we also evaluated the performance of four available tools, namely HostPhinder^13^, VirHostMatcher^14^, WIsH^14^ and DeepHost^15^. In the tests using VirHostMatcher and WIsH, the host reference database consists of all 107 bacteria genomes from NCBI; the bacteria with a sequence most similar to that of the phage sequence is taken as the prediction.

Graphage outperformed all the methods compared (Table 5). Only the AF-based model achieved performance comparable to Graphage in both accuracy and weighted F1 score; this could be due to the *k*-nearest neighbors method used in the model which generalizes well to multiclass prediction tasks. All the methods other than VirHostMatcher scored much worse in macro F1-score than in weighted F1-score, since predicting the samples from rare classes is a much more challenging task.

**Table 5.** Comparing our graph model with other models and tools on phage host species prediction tasks. The best performing model is in bold font, while the runner-up is underlined. Results are averaged over 5 random train-test splits.

| Method | Accuracy | Weighted F1 score | Macro F1 score |
| --- | --- | --- | --- |
| Graphage (Ours) | **0.940±0.001** | **0.937±0.001** | **0.730±0.024** |
| AF-based | 0.931±0.001 | 0.931±0.001 | <u>0.715±0.001</u> |
| word2vec | 0.746±0.032 | 0.725±0.033 | 0.358±0.014 |
| Sequence-based | 0.644±0.035 | 0.610±0.051 | 0.210±0.033 |
| Gapped pattern frequency-based using |  |  |  |
| *k*-Nearest Neighbors | <u>0.935±0.003</u> | <u>0.932±0.002</u> | <u>0.715±0.004</u> |
| Random Forest | 0.923±0.003 | 0.913±0.003 | 0.679±0.019 |
| MLP | 0.798±0.056 | 0.430±0.048 | 0.682±0.074 |
| ----- |  |  |  |
| HostPhinder<sup>13</sup> | 0.320±0.010 | 0.416±0.017 | 0.136±0.028 |
| VirHostMatcher<sup>14</sup> | 0.380±0.005 | 0.372±0.005 | 0.461±0.011 |
| WIsH<sup>14</sup> | 0.377±0.007 | 0.423±0.009 | 0.408±0.014 |
| DeepHost<sup>15</sup> | 0.885±0.017 | 0.870±0.023 | 0.398±0.104 |

### Necessity of GCN component

To verify the necessity of the GCN component within the GP-GCN framework, we performed an ablation study by removing the GCN while keeping other components the same. We trained new models for the four tasks and compared the performance with Graphage. The results in Supplementary Table S5 show that the models with GCN achieved superior performance on all four tasks.

For evaluation, we visually inspect the influence of GCN on the final output of Graphage on a binary classification task. More precisely, we examine the outputs of Graphage right before its final, fully-connected layers, to see how well the outputs of sequences from the same class form clusters; we compare the clusters for the case when GCN is used and when it is not. To enable visualization, we use uniform manifold approximation and projection (UMAP)^16^ to project the output values for each sequence into two dimensions. As shown in Figure 2, the outputs when GCN is used form more clearly defined clusters in accordance with their labels.

**Figure 2.**
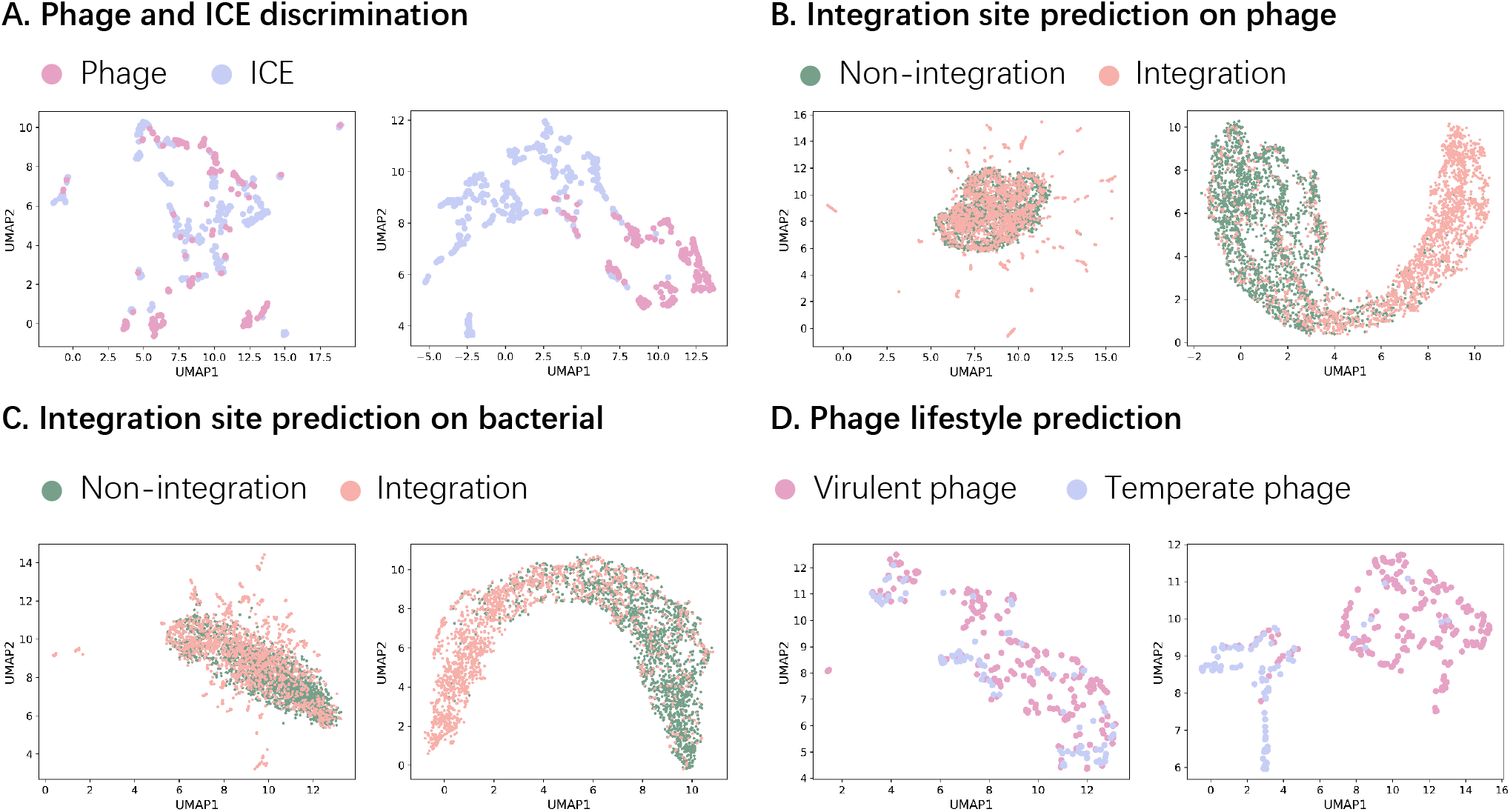
UMAP projections of the output from models with GCN and without GCN. For all subfigures A, B, C, and D, the latent features output by the models (prior to the final layer) without GCN are shown on the left, while that output by the models with GCN is shown on the right. Better separation between classes can be observed in the output of the models with GCN.

### Mining gapped patterns of more significant influence in prediction tasks

From the models of Graphage, we calculate the contribution scores for the patterns and pattern groups (see *Contribution score of gapped patterns and motifs* subsection) to mine informative patterns for the phage-related tasks.

For phage lifestyle prediction, we present the contribution score distribution for the gapped patterns in Supplementary Figure S8 A. The bimodal distribution shows that some patterns have relatively high contribution scores (0.15-0.4). In Supplementary Figure S8 B, we give the occurrence frequencies for the five gapped patterns with the highest contribution scores (left) and the frequencies for the five with the lowest contribution scores (right). We note that the gapped patterns with the highest contribution scores demonstrate significantly different frequencies between temperate and virulent phages, suggesting their significance in the classification task. We also analyze the contribution scores of the pattern groups (Supplementary Figure S8 C). The results also indicate that patterns with high contribution scores are more likely to occur in temperate phages.

For phage and ICE discrimination task and integration site prediction task, the contribution score distribution is unimodal (Supplementary Figure S9 left), with the peak <0.05, indicating that the significance of a single gapped pattern is unremarkable. However, the analyzes for the pattern groups (Supplementary Figure S9 middle and right) show that for phage and ICE discrimination, the patterns with high contribution scores are more likely to occur in ICE sequences; for integration site prediction, the patterns with high contribution scores are more likely to occur in the integration sites.

### Regulatory motifs involved in phage integration

We investigated the regulatory motifs involved to study the phage integration mechanism. First, we applied STREME^17^ to discover the motifs enriched in the sequences of the integration site with negative sequences as control. For each motif, we calculated the contribution score by removing all the possible gapped patterns in the motif from the sequences and calculating the mean absolute error between the two output probabilities. We also calculated a baseline score for each motif by removing the same number of randomly chosen gapped patterns. Then we identified the informative motifs which have higher contribution scores than their baselines. As a result, we obtained 57 informative motifs from phage integration sites and 51 motifs from bacterial integration sites. These informative motifs match with more positive sequences than the motifs with lower contribution scores than baseline (Figure 3 A and C).

**Figure 3.**
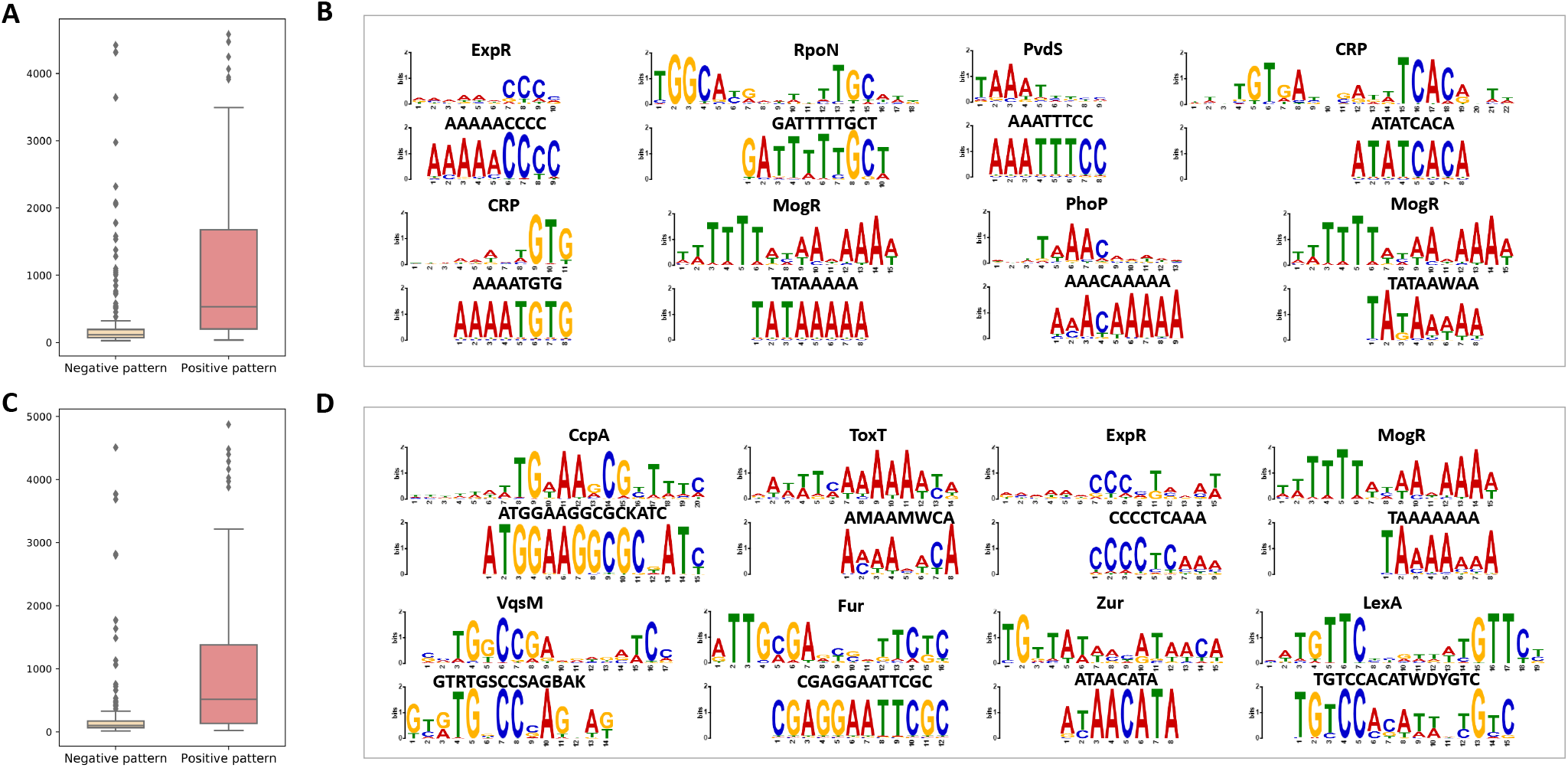
Motifs discovered in the integration site dataset match the TF-binding profiles in CollecTF. In the boxplots, we show the number of positive sequences for the enriched motifs with lower contribution scores than the baselines and higher contribution scores than the baselines in phage integration sites (A) and bacterial integration sites (C). For the motifs with higher contribution scores than baselines in phage integration sites (B) and bacterial integration site (D), we show the sequence logo alignment between the motif in CollecTF (top) and the motif in the integration site (bottom) for the top eight motifs according to the alignment *p*-value.

Next, we utilized Tomtom^18^ to compare these motifs against the CollecTF dataset^19^, which collects the transcription factor (TF)-binding sites in Bacteria. We present the sequence logo alignment for the eight motifs with the lowest alignment *p*-value (Figures 3 B and D). Specifically, for the phage integration site (Figure 3 B), the motif “GATTTTTGCT” matches with RpoN. RpoN was reported to participate in signal-response mechanisms for activities such as motility and virulence^20^. Also, the motif “ATATCACA” can be aligned to CRP, which is involved in phage replication^21^. For the bacterial integration site (Figure 3 D), the motif “ATGGAAGGCGCKATC” is consistent with CcpA, which is down-regulated after viral infection^22^. The motif “AMAAMWCA” matches ToxT, which is found in the region containing a putative integrase^23^.

### Robustness of GCN-produced embeddings

We evaluated the robustness of GCN-produced embeddings through large-scale simulated sequences. Two criteria were used to evaluate the embeddings: (1) similar sequences should have similar embeddings and vice versa, (2) embeddings should remain the same under a reasonable amount of genomic variations. As dataset we randomly selected 100 phage sequences from NCBI and simulated their genomic variants using insertion, deletion, inversion, and translocation. We generated three datasets with different variation rates and lengths (See Supplementary Method S1.2 for construction details of the simulated dataset).

To examine the first criterion, we obtained embeddings using the GCN component of the model trained for the phage and ICE discrimination task. The dissimilarity between two embeddings is computed as the Euclidean distance between the embeddings. As shown in Supplementary Figure S10, distances between the embeddings are small, even with large variations in the simulated sequences. We compared these distances to the case where the GCN is replaced with a CNN that embeds sequences into vectors of the same dimension as the GCN-produced embeddings. For all of the three datasets, GCN gave embeddings with smaller distances between the original and the mutated sequences.

### Scalability of the GP-GCN framework

Due to the very large number of biological sequences in phage analysis, any method for analyzing phage sequences must be capable of handling millions of sequences. The GP-GCN encoding framework is learning-based; after the initial training, the model can perform predictions in runtime linear to the number of sequences to infer. Additionally, each gapped pattern graph can be constructed in time polynomial to the sequence length (Supplementary Method S1.3). The computational efficiency makes the framework suitable for handling large datasets.

## Discussion

The bio-sequences are known for their frequent genetic variations, which complicates any analysis that requires homology search. Current computations of homology are mainly alignment- or AF-based. Alignment-based methods face runtime complexity issues, while AF-based methods compromise important information in the sequences. As bio-sequence databases become large and sequence diversity increases, such compromises are beginning to limit genomic analyses.

In this work, we propose a novel method for succinct biosequence representation, where each sequence is represented as a *k*-mer-encoding graph called GPG. To enable efficient comparison of these GPGs we exploit recent advances in graph neural networks to filter such graphs into low-dimensional embeddings. These ideas give rise to a framework that we call GP-GCN. Based on the framework, we developed Graphage, a phage analysis tool that currently performs four tasks: phage and ICE discrimination, phage integration site prediction, phage lifestyle prediction, and phage host prediction. Graphage achieves state-of-the-art performance compared to other methods on all four tasks.

The application of representation learning to biological sequences is not new. For instance, word2vec, a widely used natural language processing (NLP) technique^24, 25^, has been applied to obtain embeddings from sequences of the human genome^26^, as well as solve problems such as species identification^27, 28^, methyladenosine site prediction^29, 30^, and MHC binding site prediction^31, 32^. However, word2vec is based on encoding local contextual information of sequence segments and has no provision for representing global information of entire sequences. In our tests, Graphage consistently outperformed word2vec-based models.

We expect the GP-GCN framework to be useful beyond the prediction tasks currently implemented in Graphage, due to its many desirable properties, such as robustness with respect to genomic variations and high computational efficiency. Our immediate future plan is to attempt the framework on more types of sequences, e.g., RNA and protein sequences, as well as on more complex downstream tasks, such as sequence generation or sequence restoration. One application, clustering of genome sequences, is of fundamental importance to many studies such as phylogenetics, and has spurred the development of many tools^33–36^. The ability to cluster biosequences more accurately, and on larger scales, would benefit many researchers.

Other possibilities abound for the practical use of the GP-GCN framework. For instance, trained GP-GCN models encode important domain knowledge of the tasks that they were trained for. Mining gapped patterns from the models trained in the four tasks in Graphage has allowed us to uncover patterns that are possibly relevant to phage lifestyle. Another possible use of the framework is in the discovery of partial similarities between GPGs. This may be useful in situations where the GPGs are constructed from genetic materials consisting of multiple species, such as in metagenomic studies. In such a case, partial GPG similarity would suggest a similar sub-composition of the two inputs.

## Methods

### Benchmark datasets

We collected seven published datasets for the four phage-related tasks (Table 6, Supplementary Figure S11). For the task of phage and ICE discrimination we used the phage genomes from NCBI^6^ and ICE^7^ sequences from ICEBerg as the training set, following the data preparation process of GPD^8^. An independent dataset, ImmeDB^9^, including ICE and prophage sequences, was used as the test set. We applied Prokka^37^ for sequence annotation to obtain information on gene density and hypothetical protein fraction. For the task of phage integration site prediction we used a temperate phage dataset with precise boundary information^38^. We extracted the sequences of integration sites from both phage and bacterial genomes to use as positive sequences, and randomly chose sequences of the same length that are 1 *kb* away from any integration site on the phage and bacteria genomes for negative sequences. For phage lifestyle prediction, we downloaded phage sequences with empirical lifestyle data^39^ and tagged them with the lifestyle-related protein domains selected by BACPHLIP^10^. We applied Glimmer^40^ to identify the genes in phage genomes and HMMER^11^ to search for the protein domains. For phage host prediction, we combined one dataset mined from metagenomic data^8^ and one dataset mined from bacterial data^38^ with their host information. We removed the phages with unidentified hosts and kept only the taxonomies with more than 20 phage genomes. All redundant sequences were removed from each of the seven datasets.

**Table 6.**
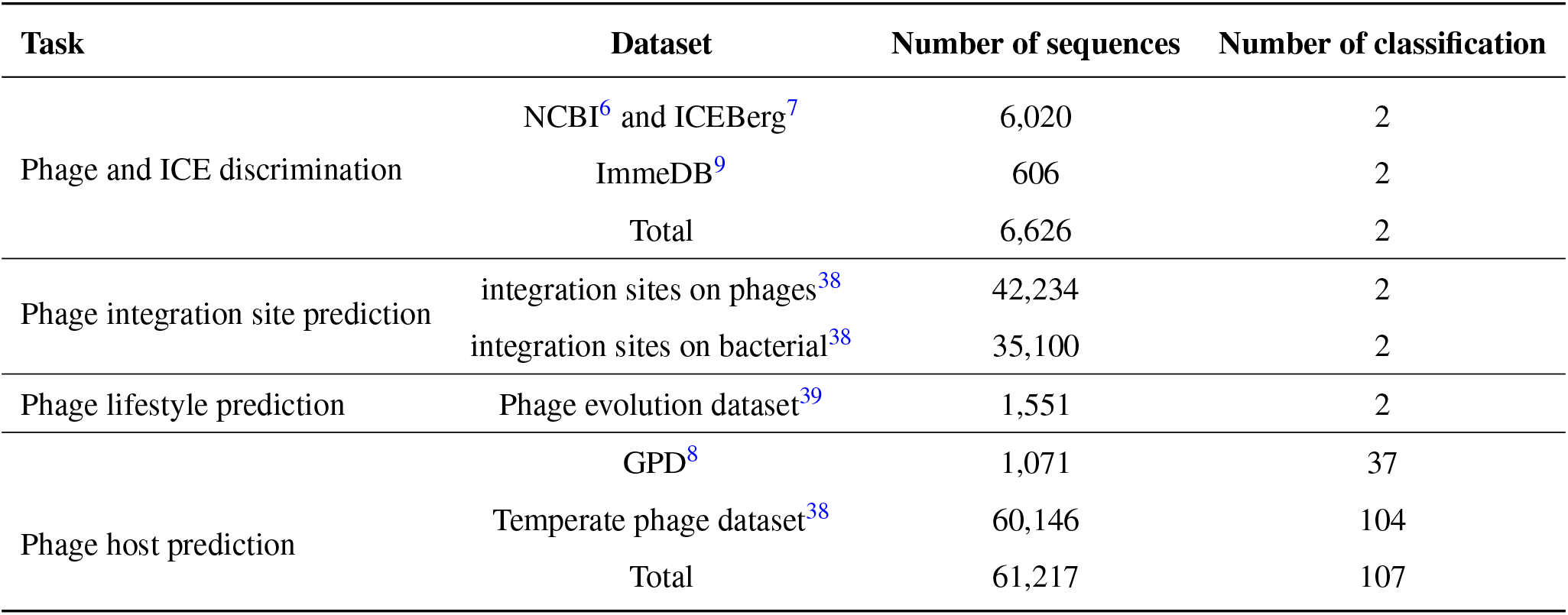
Summary of benchmark datasets in this study.

| Task | Dataset | Number of sequences | Number of classification |
| --- | --- | --- | --- |
| Phage and ICE discrimination | NCBI<sup>6</sup> and ICEBerg<sup>7</sup> | 6,020 | 2 |
|  | ImmeDB<sup>9</sup> | 606 | 2 |
|  | Total | 6,626 | 2 |
| Phage integration site prediction | integration sites on phages<sup>38</sup> | 42,234 | 2 |
|  | integration sites on bacterial<sup>38</sup> | 35,100 | 2 |
| Phage lifestyle prediction | Phage evolution dataset<sup>39</sup> | 1,551 | 2 |
| Phage host prediction | GPD<sup>8</sup> | 1,071 | 37 |
|  | Temperate phage dataset<sup>38</sup> | 60,146 | 104 |
|  | Total | 61,217 | 107 |

### Gapped pattern graph construction

#### Notation and definition

For a given sequence *S* of the alphabet {*A, G, T, C*} and a positive integer *k*, the *k*-mers of *S* are all the substrings of *S* of length *k*. Examples of 2-mers are *AA* and *GA*, while examples of 3-mers are *ACG* and *GAG*. There are a total of 4^*k*^ distinct *k*-mers for any positive integer *k*, and at most min {4^*k*^,*d* – *k* + 1} unique *k*-mers can occur in a sequence of length *d* (*d* ≥ *k*). An arbitrary indexing can be imposed over the set of all *k*-mers for any fixed *k*. Such an indexing can be used as indices in a vector, thus allowing the *k*-mer distribution of a sequence to be represented as a vector of integers.

#### Pattern graphs

Given positive integer *k*, a *pattern graph G*(*S*, *k*) of a sequence *S* is a directed graph where each vertex corresponds to a *k*-mer of *S*. For two vertices *u*, *v* ∈ *G*(*S*, *k*), an edge from *u* to *v* is in the graph if and only if the concatenation of the two *k*-mers that the vertices correspond to, *S_u_* and *S_v_* say, exist in *S*. For example, *G*(*ATGATGC*, 3) would consists of four vertices *v*_1_, *v*_2_, *v*_3_, *v*_4_ corresponding respectively to the *k*-mers *ATG, TGA, GAT*, and *TGC*; it would have exactly two edges: the edges *v*_1_ → *v*_1_ and *v*_2_ → *v*_4_. Each vertex in *G*(*S*, *k*) is assigned a number which indicates how frequently its corresponding *k*-mer occurs in *S*; similarly, each edge is assigned a number indicating how frequently the concatenation of the two *k*-mers occurs in *S*.

Two notions of frequencies are considered for the vertex and edge features: absolute or normalized counts. An absolute count of a *k*-mer is the number of times the *k*-mer occur in *S*; the normalized count of a *k*-mer is the number of times the *k*-mer occur in *S* divided by the total number of *k*-mer occurrences (i.e., *L* – *k* + 1). These variants of frequency are similarly defined for the edge features. We use normalized count for both vertex and edge features in this work. Supplementary Figure S12 shows examples of this construction.

#### Gapped pattern graphs

To allow SNPs and InDels in a sequence, we construct a *gapped pattern graph* (GPG) as follows. Let *d* be the maximum allowed gap length. The GPG *G*(*S, k, d*) is defined in the same way as the pattern graph, except for the edges. For two vertices *u*, *v* ∈ *G*(*S, k, d*), an edge from *u* to *v* is in the graph if and only if the *k*-mers that the vertices correspond to, *S_u_* and *S_v_* say, exist in *S* with *S_v_* occurring within distance *d* after *S_u_*. That is, *S_u_xS_v_* is a substring of *S*, where *x* is an arbitrary (and possibly empty) string of length at most *d*. Each edge is assigned a vector of length *d* + 1, the (*i* + 1)-th element of which indicates the frequency of *S_u_yS_v_* in *S*, where *y* is an arbitrary string of length *i*. Varying the parameter *d* allows more flexibility in the gap length, hence increasing the expressiveness of the pattern space spanned by the GPG. Examples of this construction are given in Supplementary Figure S13.

Just as every edge in a GPG can be considered a pattern of two *k*-mers with a gap of up to length *d* in between, every walk of length *l* on a GPG can be considered a pattern that consists of *l* + 1 *k*-mers, with gaps of up to length *d* in between. Given the GPG of a sequence *S*, all subsequences of *S* can be matched to one or more gapped patterns formed by such walks. Note that the other direction of this statement is not necessarily true for patterns with more than two *k*-mers.

In this work we typically let *k* = 3 and *d* = 2 (See Supplementary Tables S1-S4). However, for large values of *d*, we can compress the vector by letting some vector elements account for a range of gap lengths. For instance, we can let the first three elements indicate the frequencies of *S_u_yS_v_* where *y* has lengths of 0, 1, and 2, respectively, but let the fourth element indicate the frequencies where y has lengths of 3 or 4, the fifth element indicate the frequencies where y has lengths from 5 to 9, and so on.

#### Implementation details

GP-GCN framework uses GCN to transform each GPG into an embedding. However, current software libraries for GCP treat edges and vertices differently, whereas our formulation prefers that they be treated equally. In particular, we want edge features to be modified in the same way as vertex features.

To overcome this limitation, we insert a vertex on every edge in the GPG and assign the feature of the edge to the new vertex. The new vertex maintains the connections of the original edge; hence every edge becomes a new vertex of in-degree 1 and out-degree 1. We call these new vertices *k-mer pair vertices* to distinguish them from the *k-mer vertices* in the original GPG. This conversion does not change the information flow of the original GPG; on the other hand, all the features on the GPG are now associated with vertices, thus allowing their manipulation using standard GCN libraries.

From the construction, each *k*-mer vertex is only connected to *k*-mer pair vertices, and vice versa. An edge is set to point from a *k*-mer vertex to a *k*-mer pair vertex, then back to a *k*-mer vertex. An example of this construction is given in Supplementary Figure S14.

### Neural network architecture

After constructing the GPGs from sequences, a GCN is used to transform each GPG into an embedding. We use the PyTorch Geometric^41^ implementation of GraphSAGE^42^ for GCN (i.e., SAGEConv). Using the default settings, each graph convolutional layer computes the function,

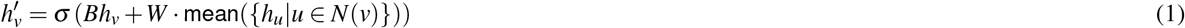

where *h_v_* is the feature of vertex *v*, *N*(*v*) is the set of all vertices in the graph where edges incident to *v* emanate from, *σ* is a non-linear activation function, mean computes the mean of the input features, and *W, B* are matrices to be learned. (Since GPGs are directed graphs, the use of such a function implies a spatial paradigm rather than a spectral one. Spectral GCN models for directed graphs exist^43–45^ but they are not yet generally available in standard libraries.)

As discussed in *Implementation Details* above, we convert edge features in the GPG into vertex features. Since the feature length of the *k*-mer vertices does not match that of the *k*-mer pair vertices in general, we use a two-step feature computation as follows. Denote a GPG as *G*(*S, k, d*) = (*V, U,E*), where *V* ={ *v*_1_, *v*_2_,…, *v*_n_} is the set of *k*-mer pair vertices, *U* = { *u*_1_, *u*_2_,…, *u*_n_} is the set of *k*-mer vertices, and *E* = {(*v, u*)|*v* ∈ *V*, *u* ∈ *U*} ∪ {(*u, v*)|*v* ∈ *V*, *u* ∈ *U*} is the set of edges. The input features for *k*-mer pair vertices *X_v_* = {*x_v_*|*v* ∈ *V*} and input features for *k*-mer vertices *X_u_* = {*x_u_*|*u* ∈ *U*} are the encoded frequencies, where 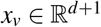 and 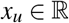. A graph convolutional layer at level *l* accepts as input a list of *k*-mer pair vertex features 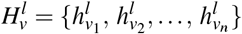 and a list of *k*-mer vertex features 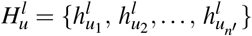. Clearly, 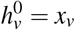 and 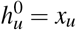. The layer outputs 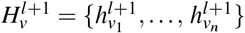 and 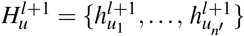 which are computed as

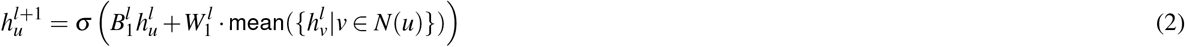

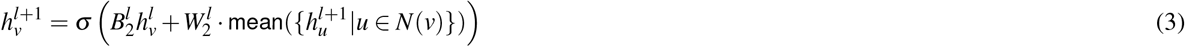

where 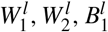 and 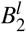 are matrices to be learned. The outputs of the graph convolutional layer at level *l* are given as inputs to the graph convolutional layer at level *l* + 1. Note that we use 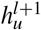 instead of 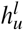 in the computation of 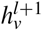 (Eqn 3) as a matter of preference.

The output of the final graph convolutional layer is concatenated and given as input to a CNN^46^ to produce the embeddings that are used in downstream tasks.

In the present work, all downstream tasks are performed with fully connected multilayer perceptron (MLP) modeled for classification. For the phage and ICE discrimination task and the phage lifestyle prediction task, we further give the genetic features to the first layer of the MLP.

### Training process and hyperparameters

We split data into training, validation, and test data (See Supplementary Table S1-S4 for details). During training, we use weighted sampling to overcome label imbalance in the training data. The sampling weight of a label is set to the inverse of its frequency in the training data. We apply mini-batch gradient descent to optimize the cross-entropy^47^ loss function

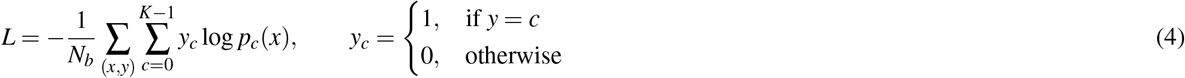

where *N_b_* is the batch size, *x* the network input, *y* the label, *K* the number of classes, and *p_c_*(*x*) the predicted probability that *x* belongs to class *c*. The Adam gradient descent algorithm^48^ is used in back-propagation with a learning rate of 10^-4^.

The neural networks are implemented in PyTorch^49^ and PyTorch Geometric^41^. Other hyperparameters and settings are summarized in Supplementary Table S1-S4. All experiments were run on an Nvidia Tesla T4 (16G).

### Contribution score of gapped patterns and motifs

As mentioned, every edge of a GPG represents a gapped *k*-mer pair pattern. Some of these gapped patterns may exert more influence in a prediction task than others. To investigate this possibility, for a given input sequence, we remove the feature of an edge from the sequence’s GPG and examine how that influences the prediction accuracy of a trained model on the sequence. More precisely, given a phage sequence *S*, we first obtain the trained output probability for the class that *S* belongs to. Then, we obtain the GPG of *S* and set the feature of a given edge in the GPG to zero, then obtain the new corresponding output probability; the mean absolute error (MAE) between the two probabilities provides us with a contribution score for the gapped pattern that corresponds to the given edge, indicating its influence on the prediction. This is repeated for every sequence and every edge of the GPG of the sequence. The total contribution of each gapped pattern is aggregated from all the differences resulting from its edge being set to zero (Supplementary Figure S15 A).

However, similar patterns are likely to influence the model simultaneously; thus, some patterns may have low contribution scores even with great influence on the model prediction. To remove this bias, we first sort the patterns in descending order according to their contribution scores. With this order, we select representative patterns, requiring a representative pattern to have Hamming distance ≥ 2 from other representative patterns. In this way, we select a distinct pattern with a high contribution score to represent a group. Next, we assign the remaining pattern to the group with a similar representative pattern (Hamming distance ≤ 1). We annotate each group with the contribution score of the representative pattern (Supplementary Figure S15 B).

For a given motif, we can also obtain the contribution score by removing all the possible gapped patterns in the motif from the sequences and calculating the MAE between the two output probabilities (Supplementary Figure S15 C). The pattern and motif scoring functions are included in the GP-GCN package.

### Competing methods

A GPG contains information of both the frequencies of *k*-mers (vertex feature) and gapped patterns of two *k*-mers (edge feature). The former corresponds to the traditional AF-based input (i.e., *k*-mer distribution), while the latter is unique to GPG. To show that the combination of these information (in the form of the GPG) is beneficial to the prediction tasks studied in this work, we compare Graphage to similar models that accept respectively (1) AF-based input, (2) word2vec-based input, (3) sequence-based input, and (4) distribution of all the gapped patterns of *k*-mer pairs. We want these input to provide at least the same amount of information available in the GPGs. Since we use GPGs of 3-mers, we use 6-mers distribution as the AF-based input, and we use 6-mers for both the word2vec- and sequence-based input. We use 3-mers for the gapped patterns of *k*-mer pairs. Below are the implementation details of the models for each of these inputs.

- The AF-based models are written using alfpy^1^. We represent each sequence with a 6-mer frequency vector, then calculate the distances between the sequences in the training set and the sequences in the test set. Prediction for a test sequence is performed by returning the label of its nearest neighbor in the training set.
- For word2vec, we apply Gensim^50^ to learn the 6-mer vector representations from the training sequences. Then a pooling strategy over the 6-mers is applied, and an optimized fully-connected neural network is used for the final prediction.
- For sequence-based methods, every sequence is converted into a vector of a fixed length. More precisely, each sequence is straightforwardly transformed into a 4 × *L* matrix with one-hot encoding (A is encoded as (1,0,0,0); C as (0,1,0,0); G as (0,0,1,0); T as (0,0,0,1)). For phage and ICE discrimination, phage lifestyle prediction, and phage host prediction, we set *L* to 50,000, which is the median length of phage sequences. Sequences shorter than 50 *kb* are padded with trailing zeros, while sequences longer than 50 *kb* are discarded. An optimized fully-connected neural network is used for the final prediction.
- Since the gapped patterns of *k*-mer pairs is a new form of input, we evaluated several conventional machine learning models in order to identify the ones that are most suitable for it. The models examined are (1) SVM (with both linear function and radial basis function kernel), (2) *k*-nearest neighbors, (3) logistic regression, (4) AdaBoost, (5) decision tree, (6) random forest, and (7) MLP. More details on the parameters used in these models are shown in Supplementary Method S1.1. We found that the best performances were achieved with *k*-nearest neighbors, random forest, and MLP, hence we only show the results with these three models.

We furthermore compared Graphage to other state-of-the-art tools as follows:

- GPD^8^ was used for comparison in phage and ICE discrimination.
- (Phage integration site prediction is a novel task with no currently available tools.)
- BACPHLIP^10^ and DeePhage^12^ were compared in phage lifestyle prediction.
- HostPhinder^13^, VirHostMatcher^14^, WIsH^14^ and DeepHost^15^ were compared in phage host prediction.

### Performance evaluation metrics

The classification performances for all four tasks considered in this work are evaluated through accuracy, F1-score, and receiver operating characteristic (ROC) curve.

The F1-score is defined as

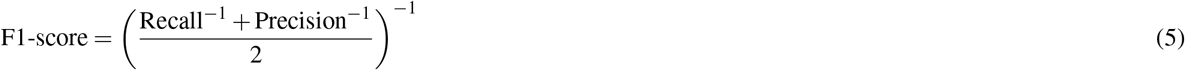

The metric is inapplicable for phage host prediction, which is a multi-class classification task. In this case, we measure weighted F1-score and macro F1-score instead.

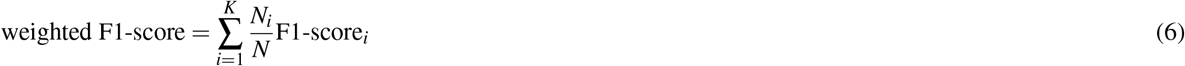

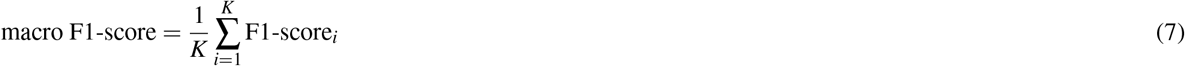

where *K* is the number of classes, *N_i_* the number of samples in class *i*, and *N* is the total number of samples.

## Acknowledgements

This work was supported by Strategic Interdisciplinary Research Grant (7020005).

## Author contributions statement

RHW designed the methods, implemented the networks, performed the experiments, and drafted the manuscript. YKN analyzed the theoretical aspects of the study and revised the manuscript. XZ revised the manuscript. JW supervised this project and revised the manuscript. SCL conceived the topic, proposed the model, and revised the manuscript. All authors have read and approved the final manuscript.

