## Supplementary materials for "Coding nucleic acid sequences with graph convolutional network"

### S1 Supplementary methods

#### S1.1 Competing methods settings

- AF-based: We applied alfpy to implement AF-based models, with word-size set to six and all other parameters set to default values (i.e., Google distance for distance metric and $k$-mer counts for vectors).
- word2vec: We applied Gensim to learn the 6-mer vector representations from the training sequences. The min_count was set to be one, and the epoch set to 20.
- Sequence-based: We used one-hot encoding to encode the sequences into matrices. The matrices were fed into a network with three convolutional layers (out_channels=64 and kernel_size=8) and three fully-connected layers (out_features=100, expect the last layer). PyTorch was used to implement the network.
- SVM (linear kernel): We applied the SVC function in sklearn package with kernel='linear', gamma=0.01, C=1, probability=True. All other parameters were set to default values.
- SVM (rbf kernel): We applied SVC function in sklearn package with kernel='rbf', gamma=0.01, C=1, probability=True. All other parameters were set to default values.
- K-Nearest Neighbors: We applied the KNeighborsClassifier function in sklearn package with n_neighbors=5. All other parameters were set to default values.
- Logistic Regression: We applied LogisticRegression function in sklearn package with penalty='l2'. All other parameters were set to default values.
- AdaBoost: We applied the AdaBoostClassifier function in the sklearn package with the default parameters.
- Decision Tree: We applied tree.DecisionTreeClassifier function in sklearn package with all parameters set to default values.
- Random Forest: We applied the RandomForestClassifier function in sklearn package with the default parameters.
- GPD: We used the scripts in the Github repository to generate feature matrices from sequences, and applied the trained model to obtain predictions.
- BACPHLIP: We applied the RandomForestClassifier function in sklearn package to reproduce the model used in BACPHLIP. All the parameters were set to the values used in the BACPHLIP paper (bootstrap=False, class_weight='balanced_subsample', min_samples_leaf=1, n_estimators=80, max_depth=40).
- DeePhage: We cut the sequence into subsequences with 1,800 bp and fed them into their 1,200-1,800 bp model. The average score for each sequence was regarded as the final prediction.
- HostPhinder: HostPhinder is a website-based tool. We used the selenium package to interact with the website automatically to submit phage sequences and obtain output.
- VirHostMatcher: We downloaded references for the 128 bacterial species from NCBI Reference Sequence Database.

- WIsH: We downloaded references for the 128 bacterial species from NCBI Reference Sequence Database. We used 6-mers and $d_2^*$ for prediction as recommended in the paper.
- DeepHost: We used our dataset to train a new model with all parameters set to their default values.

### S1.2 Simulating datasets with genetic variations

We simulated four kinds of genetic variations to evaluate the robustness of GCN-produced embeddings.

- Insertion: We randomly generated a DNA sequence from the deoxyribonucleotides dictionary {A, C, G, T} with length ranging from one to the maximum variation length. Then we randomly chose a site on the original sequence to insert the generated sequence.
- Deletion: We randomly chose a sequence ranging from one to the maximum variation length on the original sequence and removed it.
- Inversion: We randomly chose a sequence ranging from one to the maximum variation length on the original sequence and replaced it with the reverse complementary sequence.
- Translocation: We randomly chose a sequence ranging from one to the maximum variation length on the original sequence and moved it to another randomly chosen site on the original sequence.

We simulated three datasets with different variation rates and lengths. Each sequence in the first dataset contains eight variations, including two insertions, two deletions, two inversions, and two translocations. The variation lengths vary from 1 bp to 10 bp. Each sequence in the second dataset contains 20 variations, including five insertions, five deletions, five inversions, and five translocations. The variation lengths vary from 1 bp to 50 bp. Each sequence in the third dataset contains 40 variations, including ten insertions, ten deletions, ten inversions, and ten translocations. The variation lengths vary from 1 bp to 100 bp.

We collected one hundred phage sequences from NCBI as the original sequence; ten sequences were simulated with variations for each original sequence in each dataset. Thus each dataset contains one thousand sequences with variations.

### S1.3 The time complexity of pattern graph construction

**Lemma 1.** *Given a sequence $S$ of length $L$, the pattern graph $G(S, k)$ can be constructed in $O(L)$ time.*

**Lemma 2.** *Given a string $S$ of length $L$, the GPG $G(S, k, d)$ can be constructed in $O(dL)$ time.*

### S2 Supplementary figures and tables

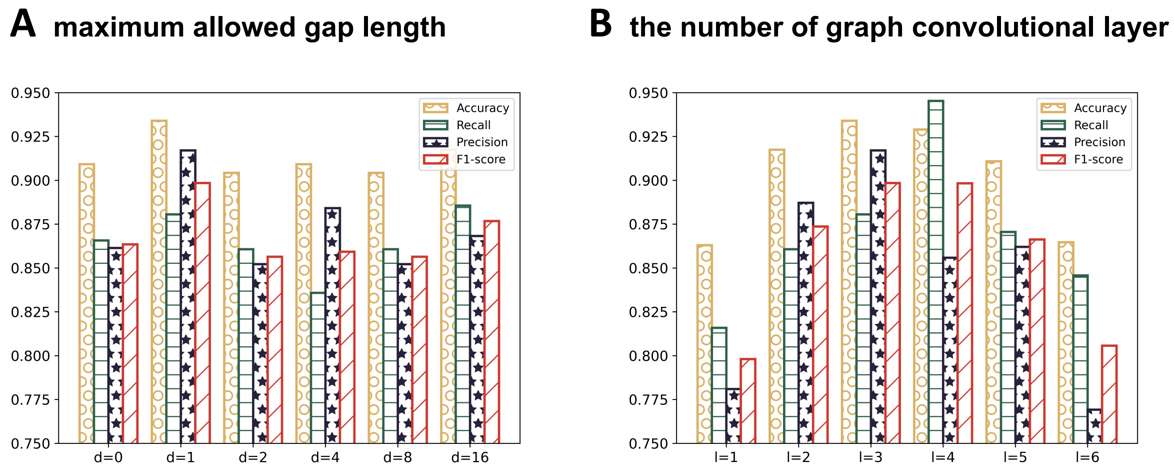


Figure S1: The effect of (A) maximum allowed gap length ($d$), and (B) the number of graph convolutional layer ($l$) on the performance of phage and ICE discrimination. (Ranges of gap lengths were used for $d$ larger than 2.)

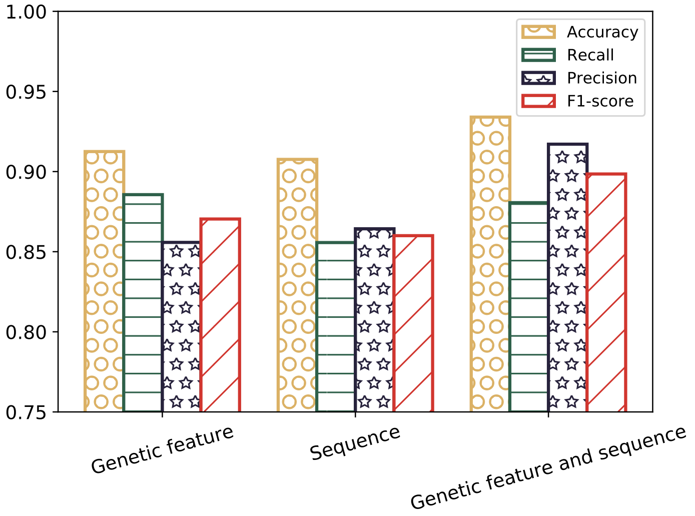


Figure S2: The phage and ICE discrimination performance of models with only sequences, only genetic features, and both sequences and genetic features as inputs.

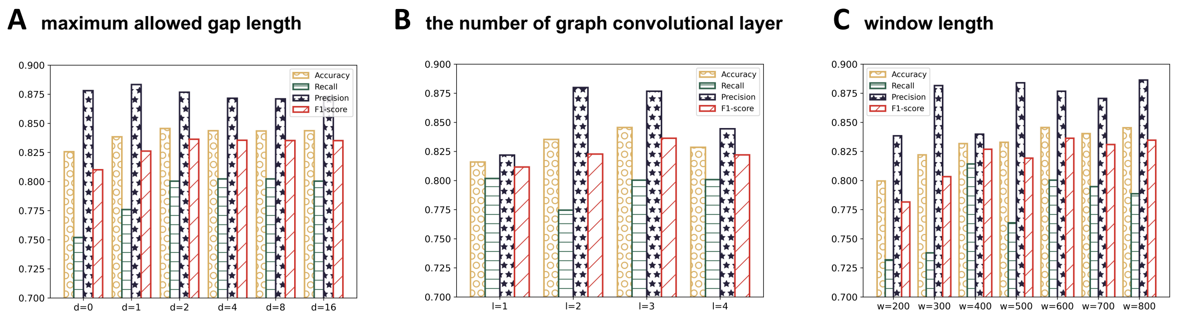


Figure S3: The effect of (A) maximum allowed gap length ($d$), (B) the number of graph convolutional layer ($l$) and (C) window length ($w$) on the performance of integration site prediction on phage genomes. (Ranges of gap lengths were used for $d$ larger than 2.)

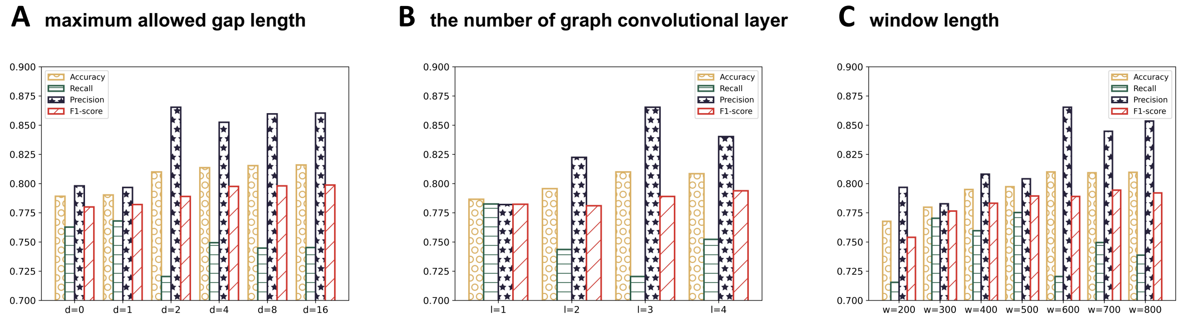


Figure S4: The effect of (A) maximum allowed gap length ($d$), (B) the number of graph convolutional layer ($l$) and (C) window length ($w$) on the performance of integration site prediction on bacteria genomes. (Ranges of gap lengths were used for $d$ larger than 2.)

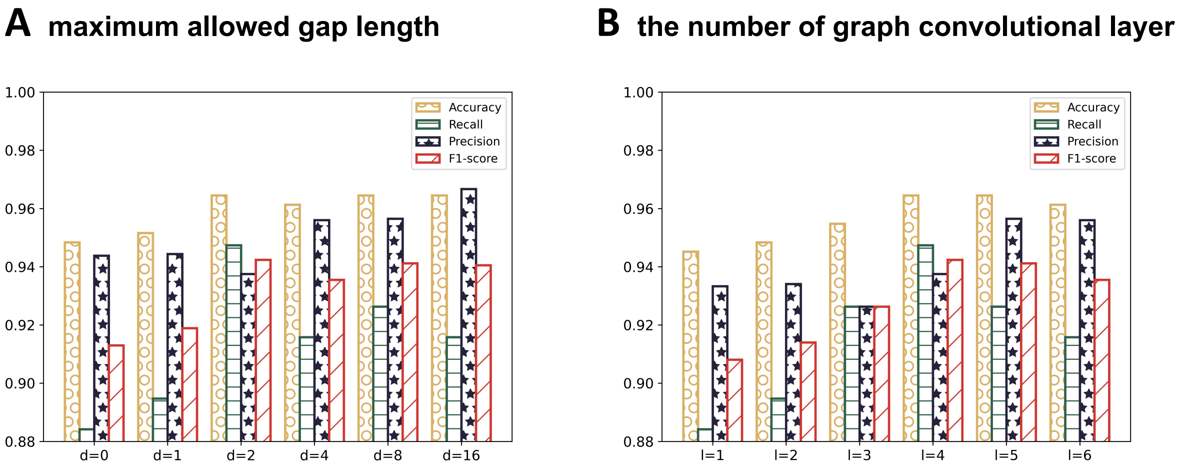


Figure S5: The effect of (A) maximum allowed gap length ($d$) and (B) the number of graph convolutional layer ($l$) on the performance of phage lifestyle prediction. (Ranges of gap lengths were used for $d$ larger than 2.)

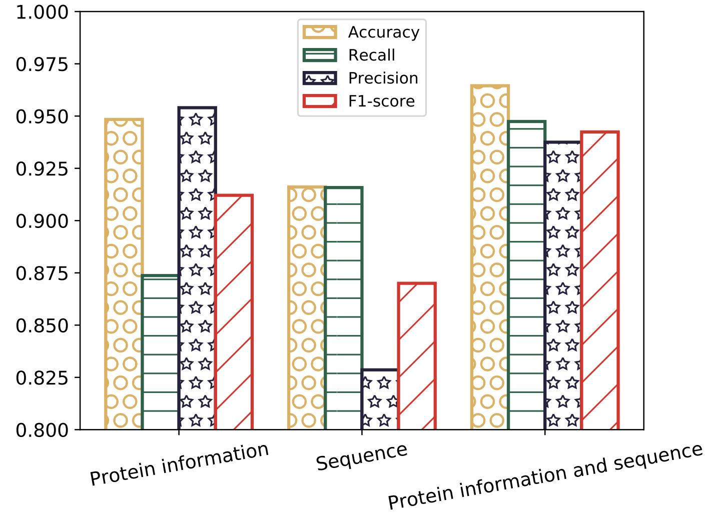


Figure S6: The lifestyle prediction performance of models with only sequences, only genetic features, and both sequences and genetic features as inputs.

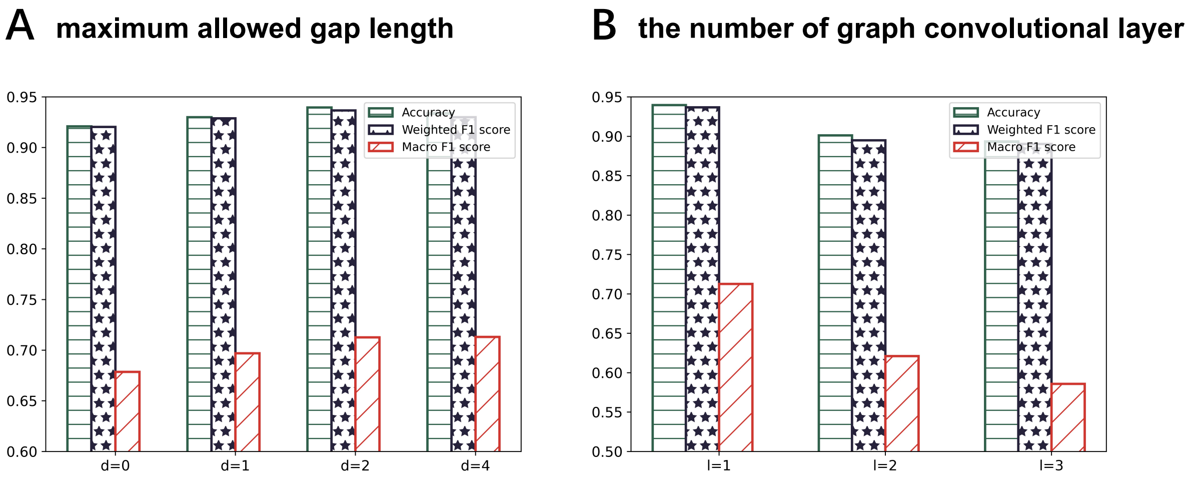


Figure S7: The effect of (A) maximum allowed gap length ($d$) and (B) the number of graph convolutional layer ($l$) on the performance of phage host prediction. (Ranges of gap lengths were used for $d$ larger than 2.)

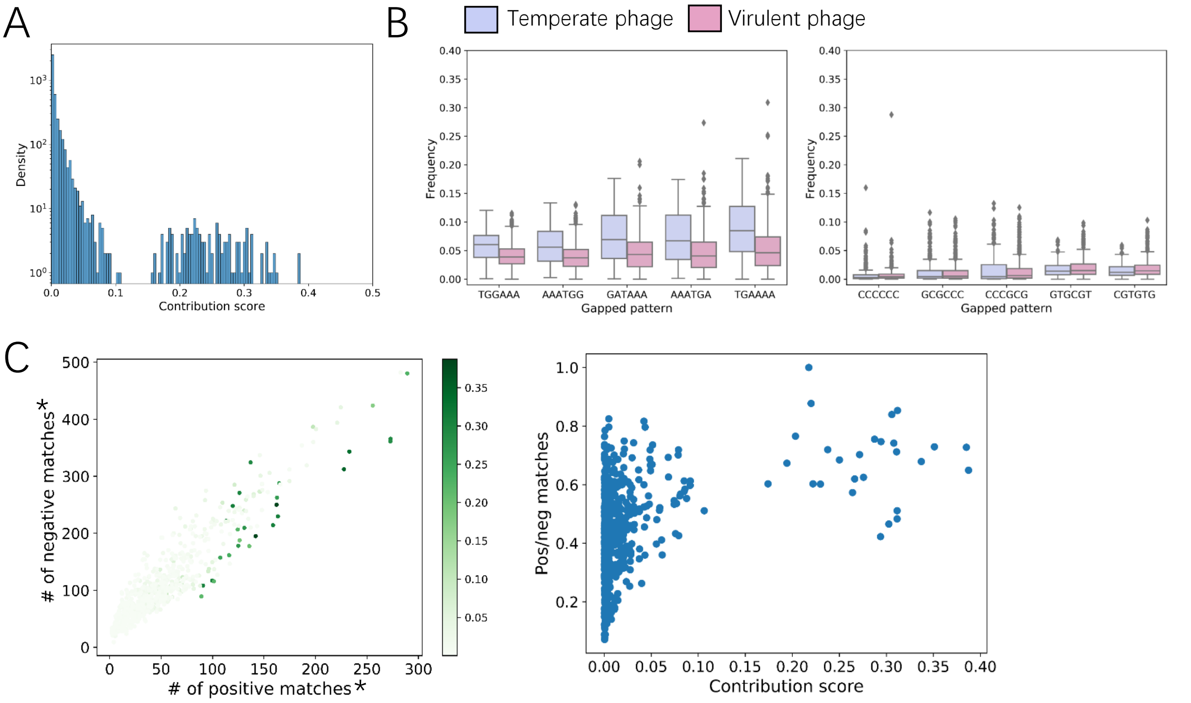


Figure S8: A. The contribution score distribution for the 4,096 gapped patterns to the phage lifestyle prediction. B. The occurrence frequencies for the five gapped patterns with the highest (left) and lowest (right) contribution scores in temperate phages and virulent phages. C. The number of positive matches and negative matches for each gapped pattern group, with the color indicating the contribution score (left). The ratio of positive matches to negative matches and contribution score for each gapped pattern group (right). *The number of positive (negative) matches is the number of positive (negative) sequences that contain the pattern. For each sequence, we only count the 10% highest matching patterns. For phage lifestyle prediction, positive sequences are temperate phages, while negative sequences are virulent phages.

| **Parameter type** | **Parameter** | **Parameter setting** | **Description** |
|---|---|---|---|
| Model parameters | label_num | 2 | The number of labels. |
| | other_feature_dim | 2 | The dimension for other features. |
| | K | 3 | The length of $k$-mer. |
| | d_n | {0,1} | The gap lengths allowed in gapped patterns. |
| | node_hidden_dim | 3 | $\|h_v^{l+1}\|$. |
| | gcn_dim | 128 | $\|h_u^{l+1}\|$. |
| | gcn_layer_num | 3 | The number of GCN layers. |
| | cnn_dim | 64 | The size of output of convolutional layers. |
| | cnn_layer_num | 3 | The number of convolutional layers. |
| | cnn_kernel_size | 8 | The kernel size of convolutional layers. |
| | fc_dim | 3 | The number of neurons for the fully connected layers. |
| | dropout_rate | 0.2 | The dropout rate. |
| | pnode_nn | Yes | Whether to embed primary features into latent space. |
| | fnode_nn | Yes | Whether to embed target features into latent space. |
| Training parameters | learning_rate | 1e-4 | The learning rate for training. |
| | batch_size | 64 | The batch size for training. |
| | epoch_n | 100 | The number of training epochs. |
| | val_split | 0.2 | The validation set size. |

Table S1: The hyperparameters used for phage and ICE discrimination in Graphage.

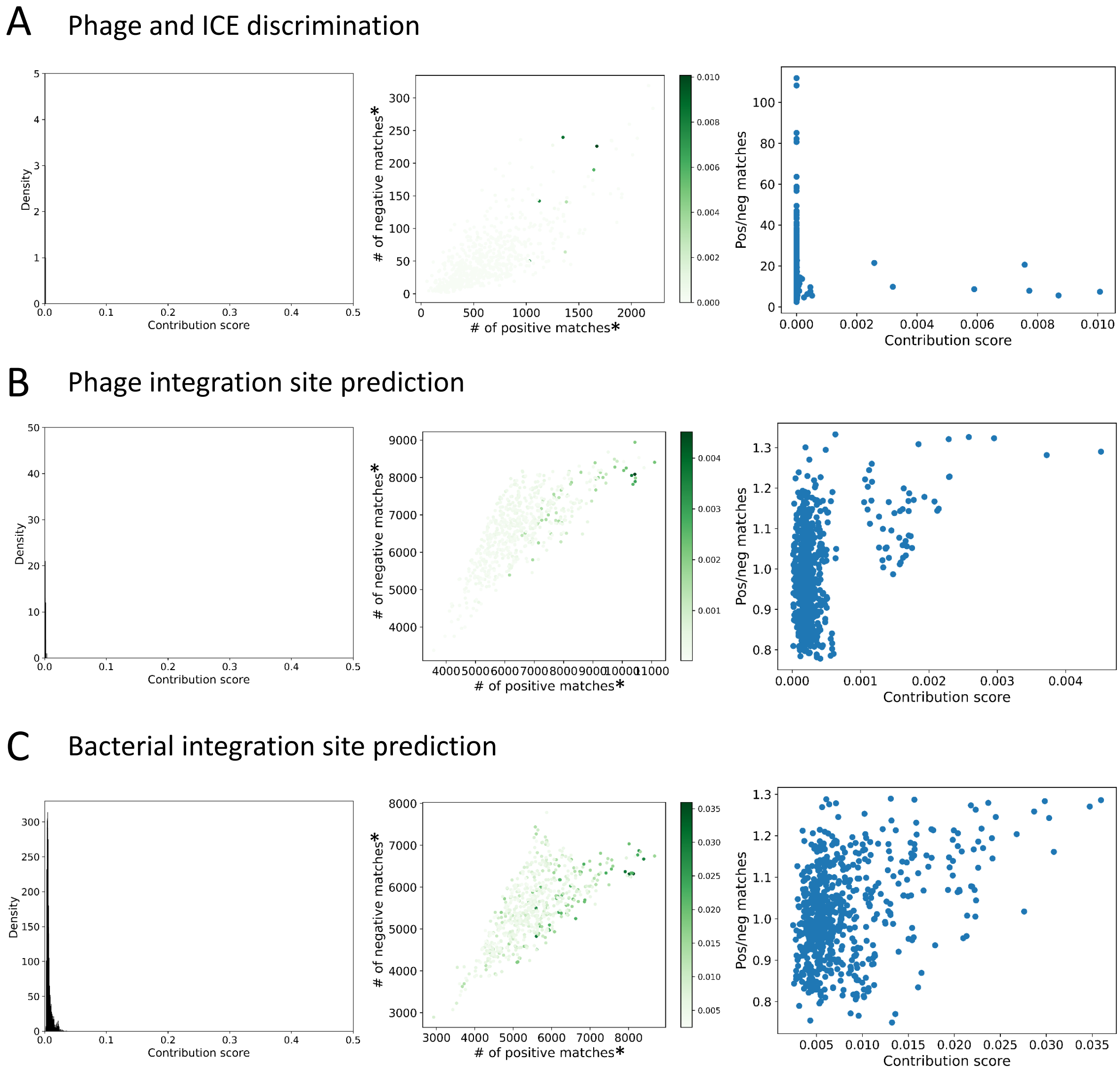


Figure S9: For A. phage and ICE discrimination, B. phage integration site prediction, and C. bacterial integration site prediction, we show the contribution score distribution for the 4,096 gapped patterns (left); For each gapped pattern group, we show the number of positive matches, the number of negative matches, and the contribution score (middle and right). *For phage and ICE discrimination, positive sequences are phages, while negative sequences are ICEs. For phage and bacterial integration site prediction, positive sequences are integration sites, while negative sequences are non-integration sites.

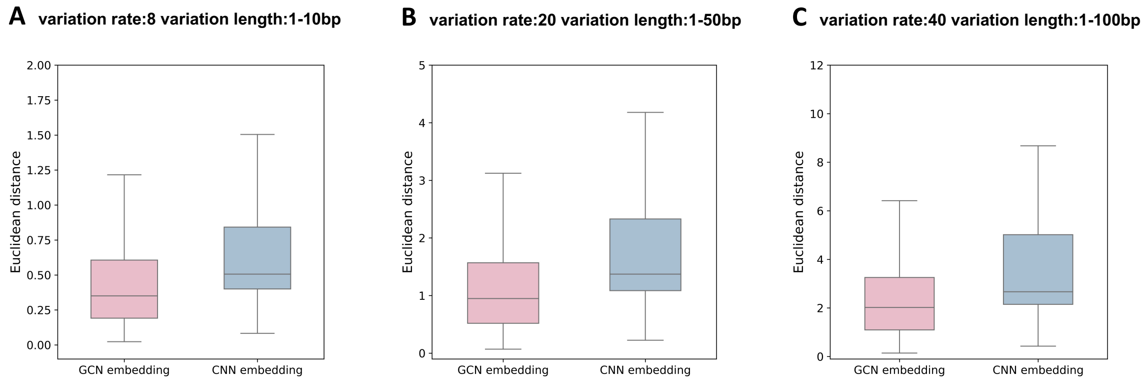


Figure S10: For the three simulated datasets, we calculated the Euclidean distance between the vectors of the original sequences and mutated sequences, from the GCN-produced embeddings and CNN-produced embeddings.

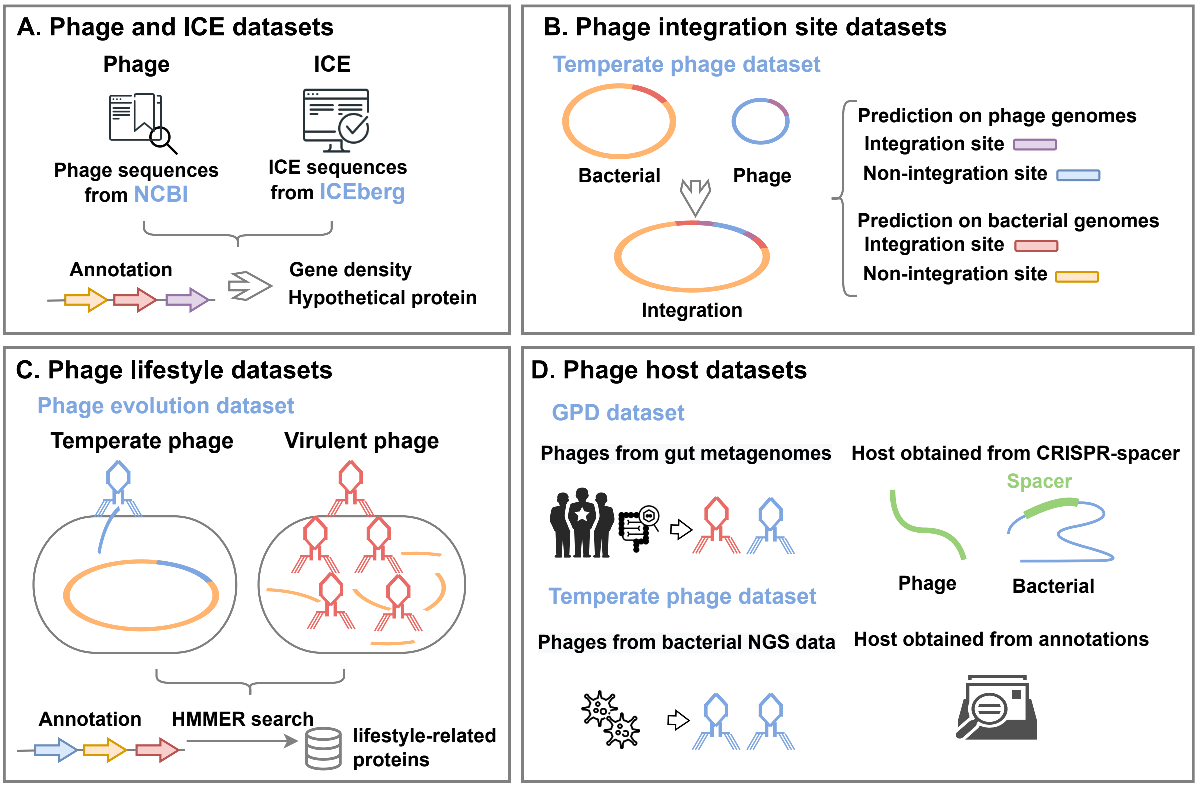


Figure S11: **Four benchmark datasets are prepared, each for a phage-related task.** A. Phage and ICE discrimination. B. Phage integration site prediction (on phage and bacterial genomes, respectively). C. Phage lifestyle prediction. D. Phage host prediction.

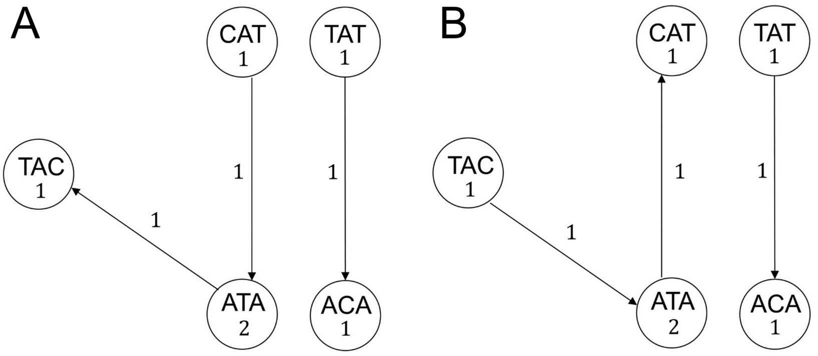


Figure S12: A. $G(\text{CATATACA}, 3)$, the 3-mer graph for the string CATATACA. B. $G(\text{TATACATA}, 3)$, the 3-mer graph for the string TATACATA. The two strings result in the same 3-mer distribution but different 3-mer graphs.

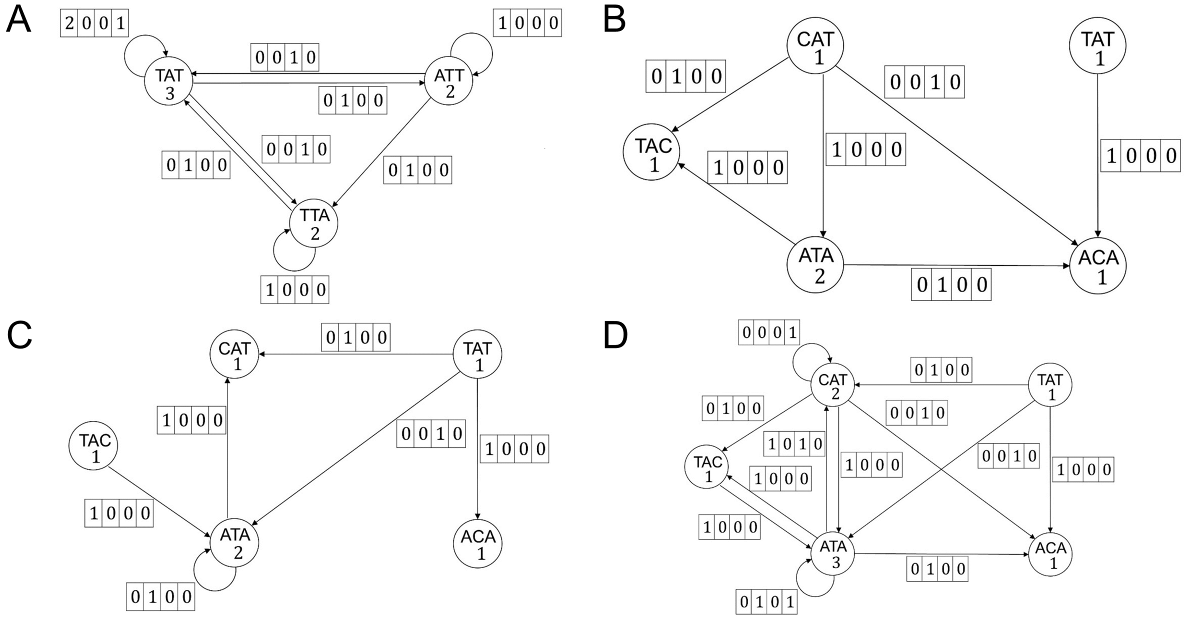


Figure S13: A. $G$(TATTATTAT,3,3), the gapped 3-mer graph for the string TATTATTAT which allows up to a gap of length 3. B. $G$(CATATACA, 3, 3), the gapped 3-mer graph for the string CATATACA which allows up to a gap of length 3. C. $G$(TATACATA, 3, 3), the gapped 3-mer graph for the string TATACATA which allows up to a gap of length 4. D. $G$(CATATACATA, 3, 3), the gapped 3-mer graph for the string CATATACATA which allows up to a gap of length 3.

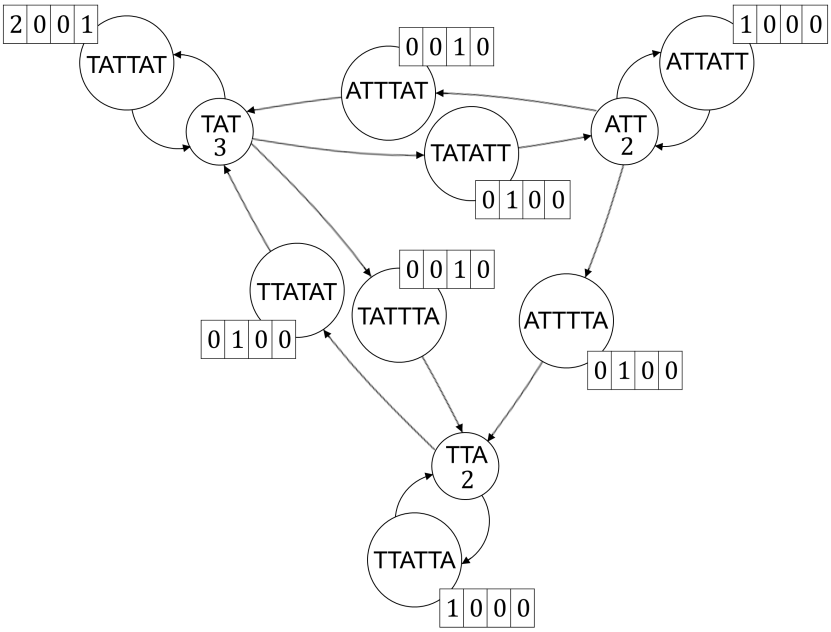


Figure S14: $S$(TATTATTAT,3,3), the gapped 3-mer graph for the string TATTATTAT which allows up to a gap of length 3, after conversion of edges into vertices. Each vertex contains a (featureless) self-loop for GCN computation, which is not shown in figure.

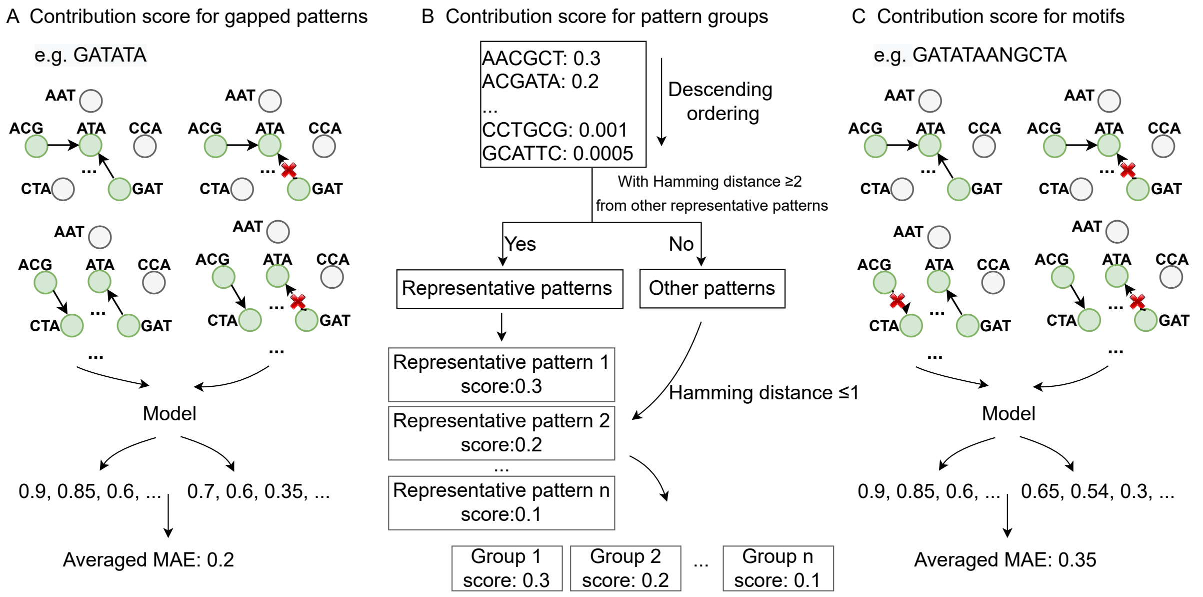


Figure S15: The calculation of contribution score for gapped patterns (A), pattern groups (B), and motifs (C).

| Parameter type | Parameter | Parameter setting | Description |
|---|---|---|---|
| Model parameters | label_num | 2 | The number of labels. |
| | other_feature_dim | 0 | The dimension for other features. |
| | K | 3 | The length of $k$-mer. |
| | d_n | {0,1,2} | The gap lengths allowed in gapped patterns. |
| | node_hidden_dim | 3 | $\|h_v^{l+1}\|$. |
| | gcn_dim | 128 | $\|h_u^{l+1}\|$. |
| | gcn_layer_num | 3 | The number of GCN layers. |
| | cnn_dim | 64 | The size of output of convolutional layers. |
| | cnn_layer_num | 3 | The number of convolutional layers. |
| | cnn_kernel_size | 8 | The kernel size of convolutional layers. |
| | fc_dim | 100 | The number of neurons for the fully connected layers. |
| | dropout_rate | 0.2 | The dropout rate. |
| | pnode_nn | Yes | Whether to embed primary features into latent space. |
| | fnode_nn | Yes | Whether to embed target features into latent space. |
| Training parameters | learning_rate | 1e-4 | The learning rate for training. |
| | batch_size | 32 | The batch size for training. |
| | epoch_n | 50 | The number of training epochs. |
| | val_split | 0.1 | The validation set size. |

Table S2: The hyperparameters used for integration site prediction in Graphage.

| Parameter type | Parameter | Parameter setting | Description |
|---|---|---|---|
| Model parameters | label_num | 2 | The number of labels. |
| | other_feature_dim | 206 | The dimension for other features. |
| | K | 3 | The length of $k$-mer. |
| | d_n | {0,1,2} | The gap lengths allowed in gapped patterns. |
| | node_hidden_dim | 3 | $\|h_v^{l+1}\|$. |
| | gcn_dim | 128 | $\|h_u^{l+1}\|$. |
| | gcn_layer_num | 4 | The number of GCN layers. |
| | cnn_dim | 64 | The size of output of convolutional layers. |
| | cnn_layer_num | 3 | The number of convolutional layers. |
| | cnn_kernel_size | 8 | The kernel size of convolutional layers. |
| | fc_dim | 100 | The number of neurons for the fully connected layers. |
| | dropout_rate | 0.2 | The dropout rate. |
| | pnode_nn | Yes | Whether to embed primary features into latent space. |
| | fnode_nn | Yes | Whether to embed target features into latent space. |
| Training parameters | learning_rate | 1e-4 | The learning rate for training. |
| | batch_size | 64 | The batch size for training. |
| | epoch_n | 200 | The number of training epochs. |
| | val_split | 0.1 | The validation set size. |

Table S3: The hyperparameters used for phage lifestyle prediction in Graphage.

| Parameter type | Parameter | Parameter setting | Description |
|---|---|---|---|
| Model parameters | label_num | 107 | The number of labels. |
| | other_feature_dim | 0 | The dimension for other features. |
| | K | 3 | The length of $k$-mer. |
| | d_n | {0,1,2} | The gap lengths allowed in gapped patterns.. |
| | node_hidden_dim | 3 | $\|h_v^{l+1}\|$. |
| | gcn_dim | 100 | $\|h_u^{l+1}\|$. |
| | gcn_layer_num | 1 | The number of GCN layers. |
| | cnn_dim | 100 | The size of output of convolutional layers. |
| | cnn_layer_num | 2 | The number of convolutional layers. |
| | cnn_kernel_size | 2 | The kernel size of convolutional layers. |
| | fc_dim | 500 | The number of neurons for the fully connected layers. |
| | dropout_rate | 0 | The dropout rate. |
| | pnode_nn | No | Whether to embed primary features into latent space. |
| | fnode_nn | Yes | Whether to embed target features into latent space. |
| Training parameters | learning_rate | 1e-4 | The learning rate for training. |
| | batch_size | 128 | The batch size for training. |
| | epoch_n | 2000 | The number of training epochs. |
| | val_split | 0.1 | The validation set size. |

Table S4: The hyperparameters used for phage host prediction in Graphage.

| Task | Model | Accuray | F1 score | AUC |
|---|---|---|---|---|
| Phage and ICE discrimination | with GCN | **0.934** | **0.899** | **0.981** |
| | without GCN | 0.769 | 0.673 | 0.789 |
| Phage integration site prediction | with GCN | **0.846** | **0.837** | **0.874** |
| | without GCN | 0.823 | 0.809 | 0.867 |
| Bacterial integration site prediction | with GCN | **0.810** | **0.789** | **0.855** |
| | without GCN | 0.789 | 0.777 | 0.848 |
| Phage lifestyle prediction | with GCN | **0.965** | **0.942** | **0.975** |
| | without GCN | 0.941 | 0.903 | 0.966 |
| | | **Accuray** | **Weighted F1 score** | **Macro F1 score** |
| Phage host prediction | with GCN | **0.940** | **0.937** | **0.713** |
| | without GCN | 0.821 | 0.787 | 0.549 |

Table S5: The performance comparison of models with and without graph convolutional layers. The best performance is bold.
